# Cortical pyramidal tract neurons constrain pathological dopaminergic motor responses

**DOI:** 10.64898/2026.10.27.758468

**Authors:** Mihai Atudorei, Gustaf Erik David Olsson, Christian del Agua Villa, Claudiu Corcea, Ulrik Gether, Maria Angela Cenci, Hartwig Roman Siebner, Mattias Rickhag

## Abstract

Pyramidal tract (PT) neurons constitute a major cortical output pathway to subcortical structures, yet their contribution to parkinsonian and dyskinetic motor states remains unclear. Here, we show that chemogenetic activation of motor cortical PT neurons selectively confines pathological dopaminergic motor responses. Unilateral PT chemogenetic activation suppressed spontaneous locomotion in healthy mice but did not improve motor deficits following an ipsilateral 6-hydroxydopamine lesion. Instead, it attenuated contraversive rotations provoked by pharmacological D1-receptor stimulation and selectively reduced axial levodopa-induced dyskinesia. Mechanistically, PT stimulation reduced acute D1-mediated c-Fos induction in the dopamine-depleted striatum, without affecting chronic ΔFosB accumulation induced by repeated levodopa exposure. These findings identify a functional interaction between cortical PT output and striatal D1-receptor signalling that limits maladaptive motor responses to dopaminergic stimulation. Our results uncover a previously unrecognised cortical mechanism regulating pathological motor behaviour and suggest PT neurons as a potential substrate for circuit-based therapies in Parkinson’s disease.

**Support or grant information:** This work was funded by a collaborative project grant “ADAptive and Precise Targeting of cortex-basal ganglia circuits in Parkinsońs Disease - ADAPT-PD” from Lundbeck Foundation to Hartwig Roman Siebner (grant nr. R336-2020-1035) and the Michael J. Fox Foundation (MJFF) for Parkinson’s Research to Mattias Rickhag (MR) (grant nr. MJFF-023112). Support was moreover received from the Danish Parkinson Association (grant nr: A816, A1266, A1601, A1966 and A2461) (MR), Bjarne Saxhofs Fond (MR), Danish Society for Neuroscience (MR), Læge Sofus Carl Emil Friis og Hustru Olga Doris Friis Legat (MR), Foundation for Neurological Research grant nr: A5600 (MR), Dagmar Marshall Foundation (MR), A.P. Møller Fonden til Lægevidenskabens Fremme grant nr: 2025-00777 (MR) and Independent Research Fund Denmark (DFF-1) grant nr: 10.46540/5283-00326B (MR).

## 1. Introduction

Motor dysfunction in Parkinson’s disease (PD) arises from abnormal activity within cortico-basal ganglia networks^1^. While the degeneration of nigrostriatal dopamine projections is a defining pathological feature, aberrant cortical activity and cortico-striatal communication are increasingly recognised as important contributors to parkinsonian motor dysfunction^2,3^. Understanding how specific cortical output pathways influence the basal ganglia may therefore provide new opportunities for circuit-based therapies.

The cerebral cortex engages the basal ganglia circuits through two major classes of pyramidal neurons: the intratelencephalic (IT) and pyramidal tract (PT) neurons^4–6^. PT neurons form a major projection system that spans over multiple motor-related structures: the striatum, subthalamic nucleus (STN), thalamus, brainstem, and spinal cord^7,8^. Their broad and lateralised connectivity places them in a strategic position to influence motor behaviour, yet their involvement in parkinsonian motor dysfunctions remains poorly understood^5^. Recent clinical studies suggest that non-invasive cortical stimulation can modulate cortico-striatal circuitry and improve motor symptoms in PD^9,10^. In parallel, rodent studies demonstrate a selective recruitment of defined cortico-fugal neural populations^11^. Together, these findings raise the possibility that PT neurons can influence pathological basal ganglia activity.

The striatal spiny projection neurons (SPNs) integrate cortical and dopaminergic inputs to regulate movement^8^. SPNs account for 95% of striatal constituents, in which conventionally, dopamine D1-receptor expressing SPNs facilitate motor output, whereas D2-receptor-expressing SPNs suppress movements^8^. As parkinsonian nigrostriatal degeneration results in striatal dopamine depletion, it elicits an imbalance between these SPNs subsets^12^. While levo-dihydroxyphenylalanine (L-DOPA) and dopamine receptor agonists alleviate this functional imbalance and can markedly improve motor function in PD^13^, chronic dopaminergic treatment gradually induces maladaptive striatal plasticity and causes debilitating L-DOPA-induced dyskinesia (LID) in the majority of patients^14,15^.

A key unresolved question is whether cortical PT output promotes or restricts dopamine-dependent motor abnormalities. Given their direct access to multiple nodes of the cortico-basal ganglia network, they are well-positioned to shape both physiological and pathological motor states^7,8^. However, their discrete impact on dopaminergic motor responses has not been directly tested.

Here, we employ a chemogenetic platform to activate genetically defined motor cortical PT neurons (i.e., Chrna2-Cre)^8,16,17^ in healthy mice and in a dopamine-depleting 6-hydroxydopamine (6-OHDA) model of PD^15,18^. We examined the motor effects of PT activation, with and without striatal D1-receptor agonism, LID, and subsequent striatal neuronal activation. Our findings reveal that PT neurons regulate motor behaviour and constrain maladaptive motor responses associated with dopaminergic stimulation, thus identifying these cortical output pathways as important modulators of pathological dopaminergic motor responses.

## 2. Methods

### 2.1 Mice

Chrna2-Cre mice were generated by the GENSAT project Tg(Chrna2-cre)OE25Gsat (Mmucd)^17^. We used hemizygous Chrna2-Cre adult mice (>12-week-old), with their corresponding wild-type (WT) littermates. The mice had *ad libitum* access to water and food, and group-housed under a 12h light/dark cycle. The experimental trials were run in accordance with the guidelines of the Danish Animal Experimentation Inspectorate (permission number: 2021-15-0201-01056), under a local animal welfare committee. All efforts were made to minimise animal suffering and to limit their numbers. Tissue samples obtained from the mice were genotyped by Transnetyx (Cordova, TN, USA) using real-time PCR. The presence of the transgene was determined using proprietary primers and probes designed by Transnetyx.

### 2.2 Experimental overview

#### 2.2.1 Naïve chemogenetic cohort

Chrna2-Cre and -WT mice (n_WT_ = 9, n_Cre_ = 10) received unilateral viral-vector injections into the motor cortex. The mice underwent open-field testing (OFT) to assess the behavioural consequences of chemogenetic activation and subsequent histological analysis or viral vector expression and activation.

#### 2.2.2 6-OHDA-lesioned dyskinetic cohort

Chrna2-Cre and -WT mice (n_WT_ = 11, n_Cre_ = 10) received a unilateral 6-OHDA infusion in the MFB and a viral-vector injection in the motor cortex. Behavioural experiments included OFT, D1-receptor agonist challenge, levodopa-induced dyskinesia (LID) assessment, and histological analyses of chronic striatal neuronal excitation.

#### 2.2.3 6-OHDA-lesioned histology cohort

Chrna2-Cre and -WT mice (n_WT_ = 4, n_Cre_ = 4) received a unilateral 6-OHDA infusion in the MFB and a viral-vector injection in the motor cortex. Following concomitant PT chemogenetic activation and dopamine D1-receptor agonism, the brains were extracted for histological analyses of acute striatal neuronal excitation.

### 2.3 Viral vector delivery and Stereotaxic surgery

All surgical interventions were conducted using a stereotaxic apparatus (Kopf Instruments, USA). During surgery, the adult mice were anesthetised with isoflurane (1.5-2%) (Attane Vet., Scanvet, Denmark). The surgical procedure was performed via a coordinate-computing device (Digital Standard, Stoelting Co., Wood Dale, ILD, USA), to which a drill (Microtorque Il #70094), a Hamilton syringe (5 µl 31 gauge, Hamilton, CA, USA) for the DREADD injection and a Neurosyringe (5 µl, 33 gauge, Hamilton, CA, USA) for the 6-OHDA lesion were connected.

#### 2.3.1 DREADD expression

AAV8-hSyn-DIO-hM3Dq-mCherry (G_q_-DREADD) (Addgene, USA), a viral vector encoding a Cre-dependent DREADD transgene - viral titer: 4.2 x 10^12^ vg/ml, was unilaterally injected into the motor cortex of Chrna2-Cre and -WT mice at the following coordinates: AP: + 1.4 mm, ML: − 1.5 mm, DV: − 0.73 mm. A total volume of 250 nl was infused at a 100 nl/min rate. Following injection, the needle was left in place for seven minutes to avoid backflow. The scalp incision was sealed with tissue adhesive (Vetbond, No. 1469SB 3M, St. Paul, MN, USA).

#### 2.3.2 Unilateral nigrostriatal depletion: 6-OHDA

6-OHDA was dissolved in 0.02% ascorbic acid (Merck/Sigma-Aldrich, Burlington, MA, USA) in isotonic saline. Using the equation:

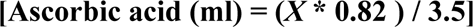

we can solve for milligrams of 6-OHDA (*X*) (Merck/Sigma-Aldrich, Burlington, MA, USA) needed to be dissolved with a chosen ascorbic acid volume. The resulting solution was kept on ice and viable to use within two hours. Due to light sensitivity, the preparation and infusion of the active 6-OHDA solution were performed under minimally illuminated conditions. A volume of 500 nl was dispensed at a flow rate of 100 nl/min into the left MFB at: AP: − 0.7 mm, ML: − 1.2 mm, DV: − 4.7 mm. The needle was subsequently left in place for seven minutes to avoid backflow.

#### 2.3.3. 6-OHDA post-operative care

Due to the severe physiological impact of the 6-OHDA lesion, the animals received intensified post-operative care for ten days. They received heating element, keeping the post-op housing cabinet at around 35 °C during the day. The weight and welfare of the animals were checked twice every day, and they were fed DietGel^®^ Boost (ClearH_2_O, Westbrook, Maine, USA), Nutella, and water-imbibed food pellets. The mice were injected with isotonic saline (no more than 1 ml/day, each injection not exceeding *bodyweight(g)/100* ml), Bupaq (100 µg/kg) (Aspen Pharma Trading Limited, Dublin, Ireland) for the first three days after surgery, and Carprofen (5 mg/kg) (Zoetis, New Jersey, USA) daily for at least a week.

### 2.4 Open-field test

The locomotion was assessed in the open-field test consisting of a squared arena divided into four quadrants of identical dimensions (50×50×35cm). A camera (USB 2.0 CMOS, Stoelting Co., IL, USA) recorded all movements via the ANY-maze 7 Video Tracking Software (Stoelting Co., IL, USA). The immobility sensitivity was set to 65% with an immobility threshold of 2000 ms. The freezing sensitivity was set to level 3 (standard), with a freezing threshold of 2000 ms. All mice were introduced into the OFT at the start of each session in a white-noise environment. The weekly starting timeslots of the behavioural stages were 9am and 12pm, respectively, over three consecutive days.

#### 2.4.1 Deschloroclozapine (DCZ) and SKF38393 treatments

##### Naïve chemogenetic cohort

DCZ (Tocris Bioscience, Bristol, UK) was dissolved in DMSO (Sigma-Aldrich, Denmark) and deionised water to a final concentration of 10 µg/mL DCZ in < 0.5% DMSO. The mice were introduced into the OFT at t = 0 min. For the first paradigm, the mice habituated for 20 minutes and then intraperitoneally injected with DCZ (100 µg/kg) (t = 20 min). For the second paradigm, the mice were intraperitoneally administered DCZ (100 µg/kg) (t = 0 min).

##### 6-OHDA-lesioned dyskinetic cohort

DCZ dihydrochloride (Hello Bio, Bristol, UK) was diluted in deionised water (5 mg/ml). For each session, the mice were left to habituate for 20 minutes for all three OFTs. All injections occurred at t = 20 min. For the first session, the mice were intraperitoneally injected with DCZ (100 µg/kg). For the second OFT, the mice were intraperitoneally injected with DCZ (100 µg/kg) and SKF38393 (3 mg/kg), and for the third OFT only with SKF38393 (3 mg/kg).

### 2.5 LID trials: Abnormal involuntary movement scoring (AIMs) and dyskinesia time scoring (DTs)

The second part of the behavioural assessment used the abnormal involuntary movement scale (AIMs)^18^ and was divided into three consecutive weeks. Axial abnormal involuntary movements are represented by contorting actions of neck and upper body towards the contralateral side of the 6-OHDA lesion. Limb abnormal involuntary movements are trembling/fluttering movements of the contralateral forelimb. Oro-facial abnormal involuntary movements are characterised by facial muscle twitching, jaw motion, and contralateral protrusion of the tongue^18^.

To provoke dyskinesia, a 1:4 solution of L-DOPA (Merck/Sigma-Aldrich, Burlington, MA, USA): benserazide decarboxylase (Merck/Sigma-Aldrich, Burlington, MA, USA) was prepared. During the first two weeks, except weekends, each mouse received a daily L-DOPA injection: 3mg/kg in the first week, 6 mg/kg in the second week^18^. During the third week, each mouse received an intraperitoneal coadministration of DCZ (100 µg/kg) with L-DOPA (6mg/kg) on the first, third, and fifth days, when the scoring was performed. On the second and fourth days, each mouse received only an L-DOPA (6mg/kg) injection. All injections and behavioural tests occurred in the same morning period from one another.

Twenty minutes after the DCZ (100 µg/kg) and L-DOPA (6mg/kg) injections, the dyskinetic time scoring (DTs) begun. DTs consists in recording the time a mouse spends while it exhibits abnormal involuntary movements. Two researchers timed axial, limb, and oro-facial dyskinesia. Each mouse was assessed for a minute, every 20 minutes. The DTs lasted for 100 minutes (five 20-minute assessment periods). The scale (AIMs) used to interpret the DTs was adapted from well-established literature^15,19^. The DTs were translated into AIMs as follows: 0 to 5 seconds of dyskinesia would give a score of 0, 6 to 24 seconds of dyskinesia would give a score of 1, 25 to 49 seconds of dyskinesia would give a score of 2, 50 to 60 seconds of dyskinesia would give a score of 3, and lastly, 50 to 60 seconds of uninterrupted dyskinesia by an external stimulus would give a score of 4.

### 2.6 Tissue preparation

We administered an intraperitoneal injection 90 minutes prior to transcardial perfusion of either DCZ (100 µg/kg) or isotonic saline (0.1 ml/g) to the naïve and dyskinetic cohorts^20^. The 6-OHDA-lesioned histology cohort (Methods 2.2.3) received a co-administration of DCZ (100 µg/kg) and SKF38393 (3mg/kg) to quantify acute neuronal excitation (Methods 2.8.3). The mice were anesthetised with a subcutaneous injection of Midazolam: Hypnorm: Saline (1:1:2). The mice were transcardially perfused with phosphate-buffered saline (PBS, 10%), followed by paraformaldehyde (PFA, 4%). The brains were extracted and fixed in a PFA solution (4%) overnight at 4 °C, then saturated in a sucrose solution (30%) for at least 48h at 4 °C. Coronal sections (30 µm) were obtained with a cryostat (Leica CM3050 S) and collected in 24-well-plates, containing antifreeze solution (11.25 mM NaH_2_PO_4_ • H_2_O, 38.25 mM Na_2_HPO_4_ •2H_2_O, glycerol in distilled water (30%)) and kept for storage at - 22 °C.

### 2.7 Immunohistochemistry

Whole-brain samples were washed three times in PBS and introduced in blocking buffer: goat serum (5%), Triton X-100 (0.3%) • PBS (10%), and bovine serum albumin (BSA, 10%). Consequently, the coronal sections were incubated overnight at 4 °C with a primary antibody in the same blocking buffer. The next day, the samples were washed twice with washing buffer (BSA (0.25%) and Triton X-100 (0.1%)• PBS (10%)). The samples were then submerged in washing buffer containing the secondary antibody for 60 min. Finally, the samples were rinsed twice with PBS (10%), mounted and air-dried on glass slides, then sealed with Prolong Gold Antifade (Thermo Fisher Scientific, USA), containing 4’,6-diamino-2-phenylindole (DAPI) to stain neuronal nuclei if needed.

### 2.8 Microscopy imaging and analysis

All imaging was performed at the Core Facility for Integrated BioImaging (CFIB) at the University of Copenhagen on the AxioScan 7 (Zeiss, Germany). The images were generated using a plan apochromat 10×/0.45 Numerical Aperture (NA) or 20×/0.8 NA M27 objective (Zeiss, Germany).

#### 2.8.1 Tyrosine hydroxylase and striatal dopamine denervation analyses

A quantitative analysis of TH immunofluorescence in the striatum was performed in both 6-OHDA cohorts (Methods 2.2.2, 2.2.3) to assess the degree of striatal dopaminergic depletion. The scans were imported to the Aligning Big Brains & Atlases (ABBA)^21^ plugin software in Fiji^22^. ABBA allows an accurate alignment of different brain compartments with the Allen Brain Atlas^23^. Consequently, our sections were exported to QuPath^24^ for analysis. TH expression of the lesioned and intact dorsal striata was measured separately, normalised against the background fluorescence of the medial septum. The difference in TH expression between the lesioned and the intact striata of all mice was calculated as an estimate of dopaminergic loss. To deem a 6-OHDA lesion successful, we set an inclusion criterion (>85%) for striatal TH loss^18^.

#### 2.8.2 Striatal ΔFosB quantification

To evaluate the simultaneous effects of chronic L-DOPA administration and Chrna2 chemogenetic activation, we quantified the cellular expression of ΔFosB - marker used for assessing chronic neuronal excitation^13^ of the 6-OHDA-lesioned dyskinetic cohort (Methods 2.2.2). Brain section images from coinciding *bregma* level (around + 1mm) underwent the same brain alignment protocol using ABBA. In QuPath^24^, a cell identification software (StarDist)^25^ allowed isolation of striatal neurons based on NeuN expression and background interference. Striatal neurons were classified as activated upon exceeding a three-fold increase in mean signal intensity to surrounding neuropile^26^. ΔFosB^+^ cell counts were displayed as cell population density (cell/mm^2^). Using an exclusion criterion, we measured the total fluorescence expression per unit area (au/mm^2^).

#### 2.8.3 Striatal and STN c-Fos quantification

The 6-OHDA-lesioned histology cohort (Methods 2.2.3) was used to study the effects in the striatum and STN of acute dopamine D1 and Chrna2^+^ co-stimulation. These were evaluated based on the expression of the acute neuronal excitation marker c-Fos^27^ from a bregma distance of approximatively + 1mm and - 2mm for the striatum and STN, respectively. An identical protocol for cellular analysis as previously described (Methods 2.8.2) was used.

#### 2.8.4 Striatal enkephalin quantification

To investigate the proportion of striatal c-Fos expression between D1- and D2-SPNs, we analysed the colocalization degree of c-Fos with the distinct D2-SPN marker enkephalin^28^. Sections from a bregma distance of approximatively + 1mm, from the 6-OHDA-lesioned histology cohort (Methods 2.2.3) were stained for enkephalin, c-Fos, and NeuN. Two same-sized regions of interest (ROIs) from the dorso-medial and dorso-lateral striata were selected. For each ROI, the c-Fos and NeuN signals were quantified following the same protocol for cellular analysis (Methods 2.8.2), while enkephalin-expressing cells were manually quantified.

### 2.8 Statistical analysis

The locomotor metrics from the OFT (distance travelled, time freezing, time immobile, rotations and rotational bias) were analysed either through a t-test for column graphs, or through a two-way repeated measure analysis of variance (ANOVA). For the post-hoc analysis, a Šídák’s multiple comparison analysis was conducted. For the neuronal quantification, a one-way repeated measure ANOVA was used to compare the means among more than two groups based on an individual independent variable. Here, for post-hoc analysis, Tukey’s or Šídák’s multiple comparisons tests were conducted. All results are presented as mean ± standard error of the mean (SEM). A significance level of *p < 0.05 was fixed for all statistical analyses. All analyses were performed in Graph Pad Prism 11.0.2.92.

## 3. Results

### 3.1 Chrna2-Cre mice provide selective access to cortico-fugal PT neurons

Based on available data from the GENSAT project^17^ and from studies on the repartition of the cortico-striatal system^4,8^ the Chrna2-cre mouse line mimics a somatic PT neuronal expression, with cortifugal efferents to various projection areas^1^ (Fig. 1c). To achieve region- and cell type-specific neuronal transduction, Cre-dependent AAV coding for an mCherry-tagged excitatory DREADD (hM3Dq-mCherry) was injected in the motor cortex in Chrna2-Cre and -WT mice^6,7^ (Fig. 1a,b). Consistent with previous characterization of the Chrna2-Cre line, mCherry expression was restricted to layer 5 PT cortico-fugal neurons and absent in wild-type littermates. Whole-brain analysis revealed robust unilateral transduction of Chrna2^+^ neurons and extensive axonal projections throughout motor-related brain regions. Dense fibre innervation was observed in the dorsolateral striatum, with additional projections detected in the thalamus, STN, substantia nigra, ventral tegmental area, cerebral peduncle, and nucleus of Darkschewitsch (Fig. 1c). These findings confirm that the targeted neuronal population forms part of a widespread cortico-fugal motor network.

**Fig. 1:**
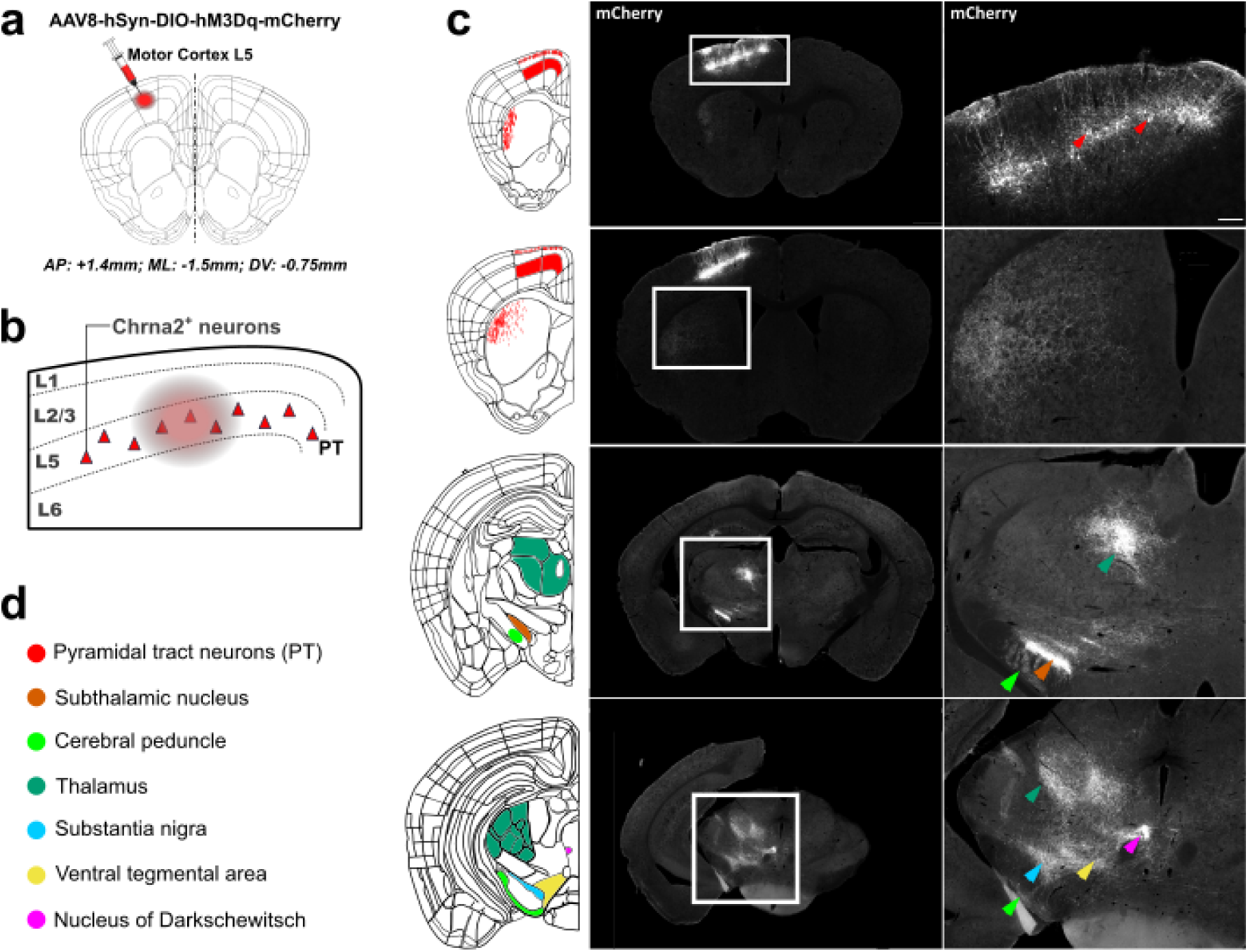
Transduction maps of Chrna2+ neurons and their axonal afferents to deeper brain regions. **a** Name of the DREADD construct (AAV8-hSyn-DIO-hM3Dq-mCherry) delivered in the mouse motor cortex layer 5 and depiction of the exact injection site with coordinates; AP: anterior-posterior, ML: medial-lateral, DV: dorso-ventral. **b** Schematic representing the somatic location of transduced PT neurons (Chrna2^+^ neurons, red) within the motor cortex and their intralayer repartition. The injection location is highlighted by a red disk. **c** Endogenous, non-antibody-amplified, representative Chrna2-Cre coronal brain slice, depicting Chrna2^+^ transduced somata and whole-brain axonal innervation (white) in coronal brain slices; approximate *bregma* levels from top row to bottom row: AP = + 1.1 mm, AP = + 0.5 mm, AP = - 1.9 mm, and AP = - 2.9 mm, respectively. Overview scale bar: 1000 µm (left column), zoomed-in scale bar: 200 µm (right column). **d** Locations of corresponding colours where positive Chrna2^+^ signal was detected. All coronal whole-brain schematics in this figure were adapted from the Allen Brain Atlas platform. Coronal slices thickness: 30 µm, AP = + 1mm.

### 3.2 Chemogenetic activation of Chrna2^+^ PT neurons suppressed spontaneous locomotor activity

The chemogenetics platform enables a tonic activation of discrete neuronal subsets to delimit their role and functional connectivity in various behavioural assessments using exclusive ligands such as DCZ^20,29–31^. Inspired by the behavioural assessment previously performed by our group^20^, we evaluated the motor effects resulting from chemogenetic activation of Chrna2^+^ neurons in the OFT. Across two independent stimulation paradigms, chemogenetic activation consistently reduced motor output. Observations following the first paradigm with DCZ administration (100 µg/kg) at t = 20 min revealed a reduction in total distance travelled (p = 0.0332) and total rotations (p = 0.0309) compared with wild-type controls (Fig. 2c,f). In the second paradigm, administration of DCZ at t = 0 min led to a similar effect resulting in reduced distance travelled (p = 0.0200), fewer rotations (p = 0.0432), and increased immobility (p = 0.0458) (Fig. 2i,j,l). In contrast, neither paradigm affected freezing behaviour or rotational bias (Fig. 2b,e,h,k). Thus, activation of Chrna2^+^ PT neurons produced a robust motor-suppressive phenotype, characterised by reduced overall locomotion and increased immobility, without inducing lateralised behaviour.

**Fig. 2:**
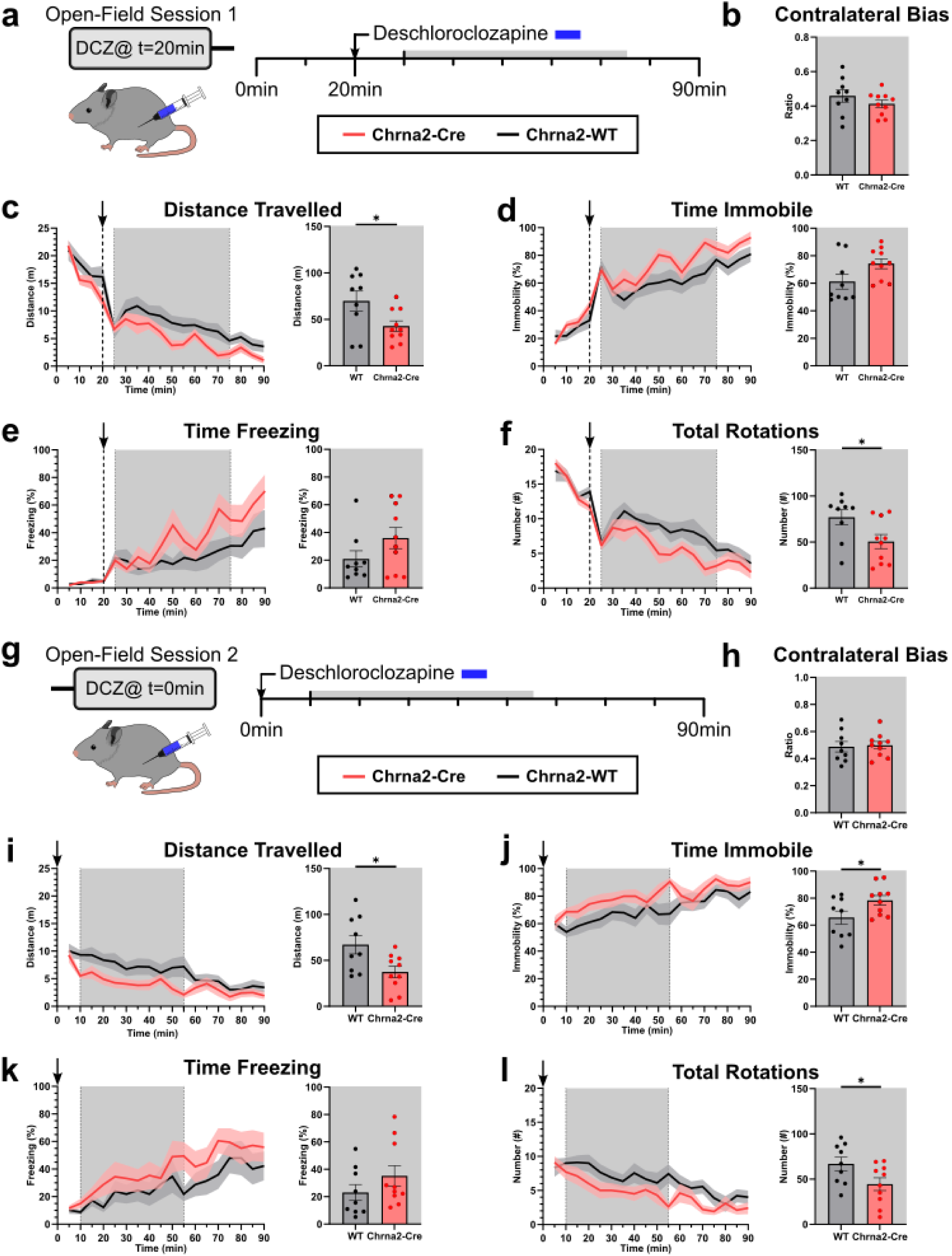
Chemogenetic activation of Chrna2+ neurons has a motor-inhibitory effect. **a** Experimental timeline of the first open-field test session. DCZ (100 µg/kg) was intraperitoneally administered at t = 20 min. **b** Contralateral bias column graph: p = 0.2765. **c** Distance travelled: genotype, F_(1,17)_ = 5.372, p = 0.0332; time F_(8, 136)_ = 9.983, p < 0.0001; interaction, F_(8,136)_ = 0.5720, p = 0.7995. **d** Percentage of time spent immobile: genotype, F_(1,17)_ = 4.091, p = 0.0591; time F_(8, 136)_ = 8.966, p < 0.0001; interaction, F_(8,136)_ = 1.016, p = 0.4266. **e** Percentage of time spent freezing: genotype, F_(1,17)_ = 2.381, p = 0.1413; time F_(8, 136)_ = 5.421, p < 0.0001; interaction, F_(8,136)_ = 1.443, p = 0.1842. **f** Total rotations: genotype, F_(1,17)_ = 5.537, p = 0.0309; time F_(8, 136)_ = 8.971, p < 0.0001; interaction, F_(8,136)_ = 0.7536, p = 0.6441. **g** Experimental timeline of the second open-field test session. DCZ (100 µg/kg) was intraperitoneally administered at the start of the paradigm (t = 0 min). **h** Contralateral bias column graph: p = 0.8115. **i** Distance travelled: genotype, F_(1,17)_ = 6.594, p = 0.0200; time F_(8, 136)_ = 2.068, p = 0.0431; interaction, F_(8,136)_ = 0.6737, p = 0.7140. **j** Percentage of time spent immobile: genotype, F_(1,17)_ = 4.642, p = 0.0458; time F_(8, 136)_ = 2.208, p = 0.0304; interaction, F_(8,136)_ = 0.9158, p = 0.5055. **k** Percentage of time spent freezing: genotype, F_(1,17)_ = 1.612, p = 0.2213; time F_(8, 136)_ = 2.749, p = 0.0076; interaction, F_(8,136)_ = 0.7639, p = 0.6351. **l** Total rotations: genotype, F_(1,17)_ = 4.772, p = 0.0432; time F_(8, 136)_ = 2.556, p = 0.0126; interaction, F_(8,136)_ = 0.6904, p = 0.6994. The dashed line and/or arrow on all time plots depict the moment the DCZ injection occurred. The column graphs (**b**-**f**) are based on the TOI = 30 – 75 min, the column graphs (**h**-**l**) are based on the TOI = 10 – 55 min. The first ten minutes post-injection(s) were discarded to avoid unanticipated auditory or olfactory disruptions. Statistical analysis: RM two-way ANOVA for the time plots and unpaired student’s t-test for the column graphs, *p < 0.05, Chrna2-Cre vs. Chrna2-WT. All results are shown as mean ± SEM. n_Chrna2-Cre_ = 10, n_Chrna2-WT_ = 9.

### 3.3 DCZ induced a robust activation of Chrna2^+^ neurons

To validate if our DREADD construct (AAV8-hSyn-DIO-hM3Dq-mCherry) successfully transduced Chrna2 neurons and if these were activated by DCZ, a histological assessment of mCherry and c-Fos signals was performed. Immunohistochemical evaluation exclusively revealed extensive mCherry expression in the Chrna2-Cre group (Fig. 3). Furthermore, the Chrna2-Cre mice injected with DCZ revealed a strong c-Fos expression, colocalizing with mCherry-positive Chrna2^+^ neurons, whereas this effect was absent in Chrna2-WT and saline-treated Chrna2-Cre mice (Fig. 3). These results confirmed an efficient transduction and activation of Chrna2^+^ neurons, linking their stimulation to the behavioural effects induced by DCZ (100 µg/kg) (Fig. 2).

**Fig. 3:**
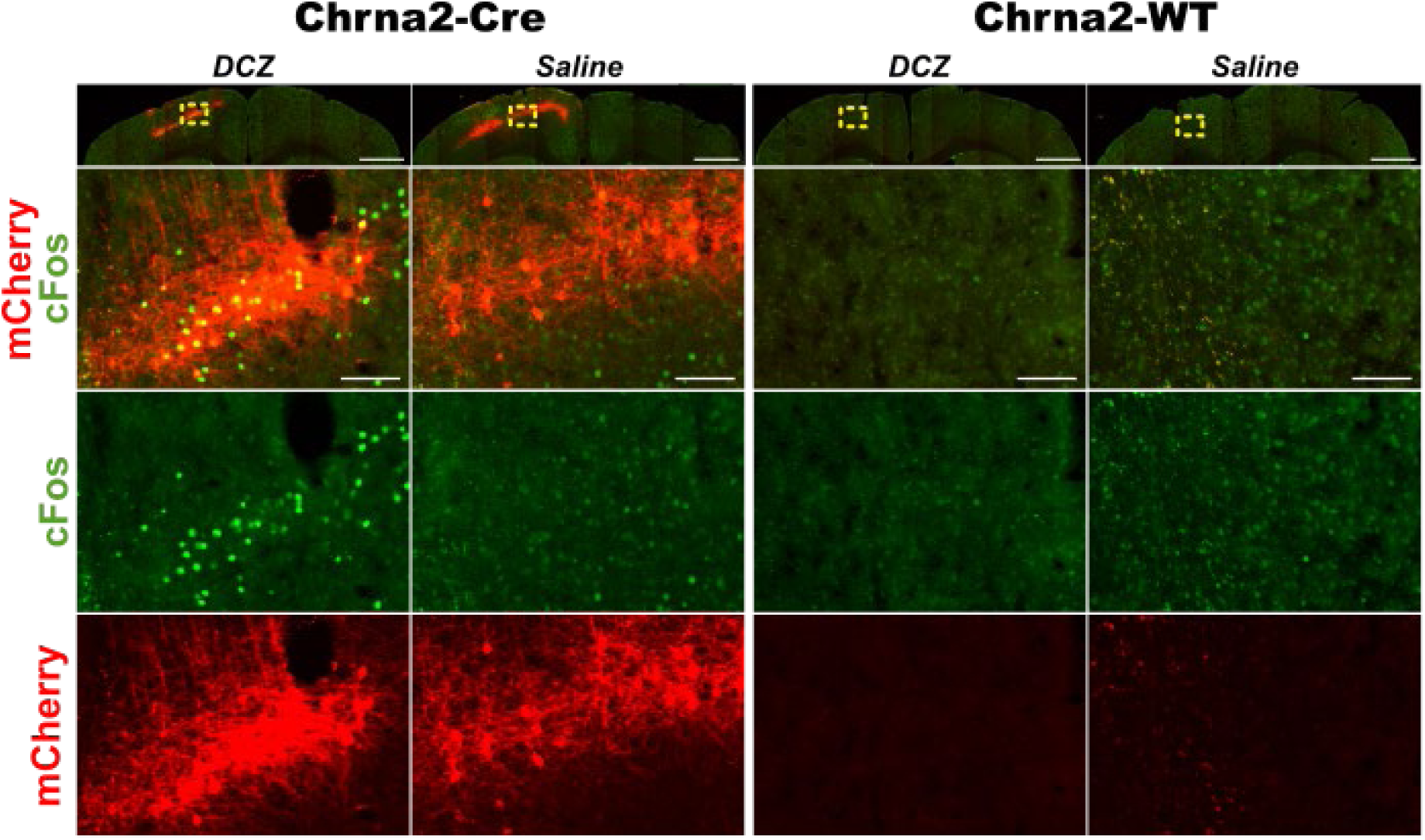
Immunohistochemical validation of Chrna2+ neuronal activation and cortical DREADD transduction map. First row: representative fluorescent images of the motor cortex in AAV8-hSyn-DIO-hM3Dq-mCherry -infused Chrna2-Cre and -WT animals challenged with 100 µg/kg DCZ or 0.01 ml/g saline. Each column is a zoomed-in ROI of the respective first row individual, highlighting Chrna2 neuronal transduction (mCherry, red) and/or neural activation (c-Fos, green) Overview scale bar: 1000 µm, zoomed-in scale bar: 100 µm. Coronal slices thickness: 30 µm, AP = + 1mm.

### 3.4 Chemogenetic experimental outline and validation of the 6-OHDA lesion model

Prompted by the results from chemogenetic stimulations in intact Chrna2-Cre mice, we applied an existent behavioural assessment protocol to a hemiparkinsonian mouse model^32^. The mice were unilaterally injected with 6-OHDA in the MFB^15^ in conjunction with the AAV-DREADD surgery (Fig. 4a). To determine whether PT neuronal activation modulates motor dysfunctions in parkinsonian conditions, we combined this activation with open-field testing and pharmacology (Fig. 4b); and subsequently with AIM/DT scorings of dyskinetic behaviours upon a chronic levodopa regimen^18^ (Fig. 4c). Successful chemogenetic targeting was confirmed by mCherry and c-Fos co-localization following DCZ administration (Fig. 4d). All animals exceeded the predefined inclusion criterion (>85%) of striatal dopamine depletion^18^ (Fig. 4e,f), confirming successful lesion induction.

**Fig. 4:**
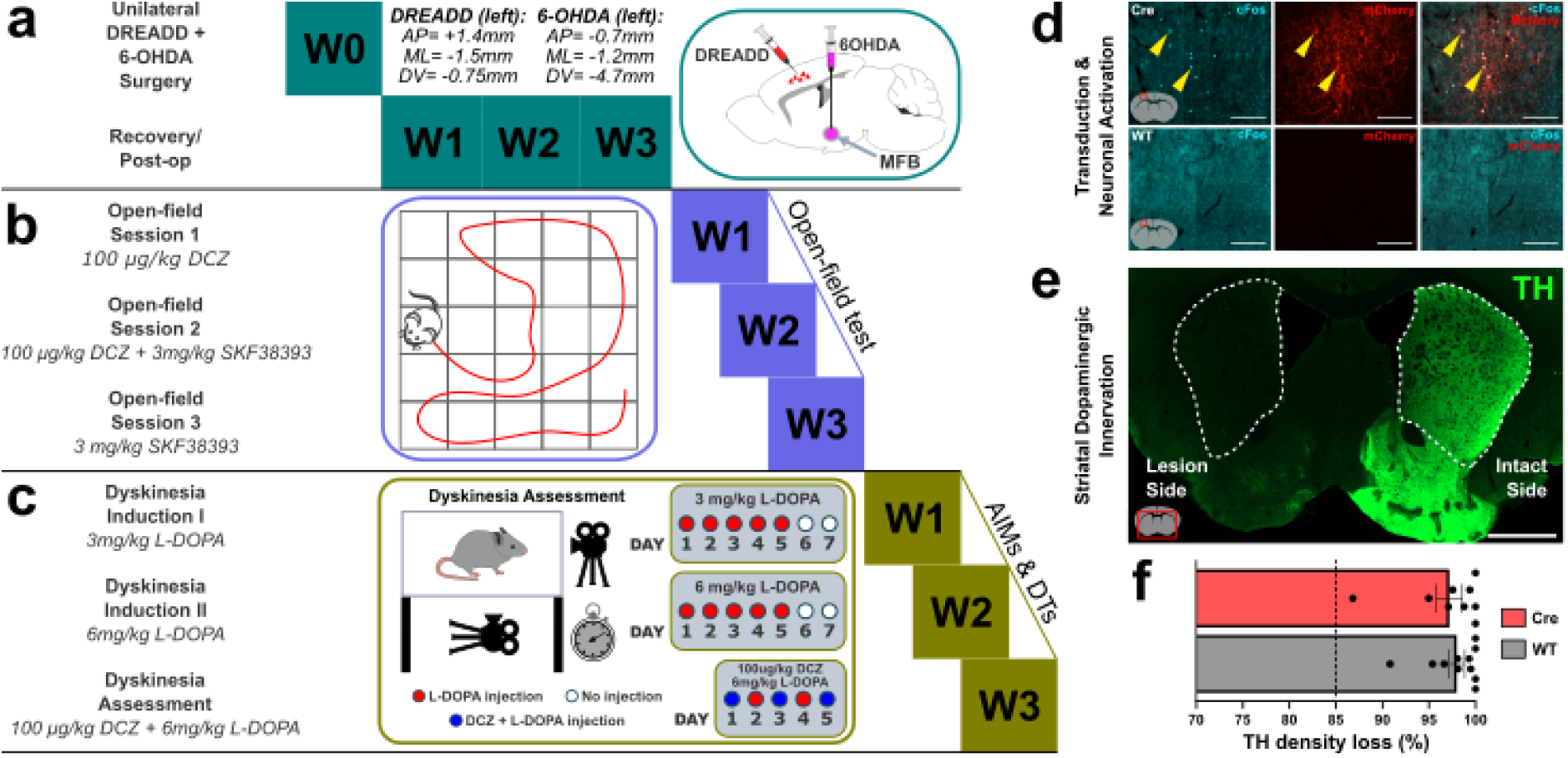
Experimental design, dopaminergic lesion validation, and cortical Chrna2+ DREADD transduction and activation. **a** Surgical procedure (one week) and post-operational period (three weeks). Each mouse was given a unilateral injection of 500 nl 6-OHDA into the MFB along with an ipsilateral injection of 250 nl excitatory DREADD (AAV8-hSyn-DIO-hM3Dq-mCherry) into the M1 motor cortex. **b** Open-field testing paradigms and schematic depicting the behavioural arena. Following the three-week post-op period, the mice underwent three separate open-field tests, each one week apart. The animals were challenged with 100 µg/kg DCZ and/or 3 mg/kg SKF38393. **c** Dyskinesia assessment experimental design. Five days a week, for three consecutive weeks, the mice were injected with 3 mg/kg L-DOPA in the first week and with 6 mg/kg L-DOPA in the second week. During the third week, the mice received 6 mg/kg L-DOPA with 100 µg/kg DCZ on day 1, day 3, and day 5, while receiving only 6 mg/kg L-DOPA on day 2 and day 4. The schematic represents how the animals were recorded and timed during LID to generate AIMs and DTs. **d** Representative cortical images of Chrna2^+^ transduction (mCherry, red) and neural activation (c-Fos, blue) between a Chrna2-Cre mouse (top row) and a Chrna2-WT mouse (bottom row). Scale bar: 200 µm. **e** Representative image of striatal TH repartition (green) in a coronal brain section of a 6-OHDA-lesioned mouse depicting the TH density difference between the intact striatum and the denervated one. Scale bar: 1000 µm. **f** Quantification of TH density loss in the dorsal striatum. The data are expressed as % of TH density loss in the lesioned striatum compared to the intact striatum between Chrna2-Cre (red) and Chrna2-WT (grey). A dashed line at 85% TH density loss serves as an inclusion threshold to a successful unilateral 6-OHDA lesion. Note that all mice were above threshold. The results are shown as mean ± SEM. n_Chrna2-Cre_ = 9, n_Chrna2-WT_ = 11. (W = Week).

### 3.5 Chrna2^+^ activation inhibited D1-receptor-mediated rotations

We wanted to investigate how motor output manifested in the open-field upon PT activation, with or without D1 agonism, in a 6-OHDA lesion model. Ipsilateral rotations are well-documented phenotypes of this neurotoxin model^33–36^. Three subsequent OFTs were performed in which animals would receive DCZ (100 µg/kg) (Fig. 5a), DCZ (100 µg/kg) and SKF38393 (3 mg/kg) (Fig. 5c) or SKF38393 (3mg/kg) (Fig. 5e) at t = 20 min. As opposed to the naïve mice resulting phenotype (Fig. 2), Chrna2^+^ activation during severe dopamine depletion exhibited no effects on distance travelled (p = 0.3325), rotations (p = 0.2738), contralateral (p = 0.7070) or ipsilateral (p = 0.2701) rotations (Fig. 5b). To pharmacologically provoke hyperlocomotion and contralateral rotating behaviours, we opted to administer the dopamine D1 partial agonist SKF38393^18,34,37^ in the second open-field paradigm, concomitantly with DCZ (100 µg/kg). In this setup, activation of Chrna2^+^ neurons resulted in significantly reduced total rotational behaviour (p = 0.0276) and decreased contralateral rotations (p = 0.0287), compared with WT controls (Fig. 5c,d). No changes were observed in overall locomotion (p = 0.1738) or ipsilateral rotations (p = 0.3806). Interestingly, subsequent SKF38393 (3 mg/kg) administration alone also resulted in reduced total rotations (p = 0.0212) and contralateral rotations (p = 0.0194) (Fig. 5e,f), while overall locomotion (p = 0.5499) and ipsilateral rotations (p = 0.0737) remained unaffected.

**Fig. 5:**
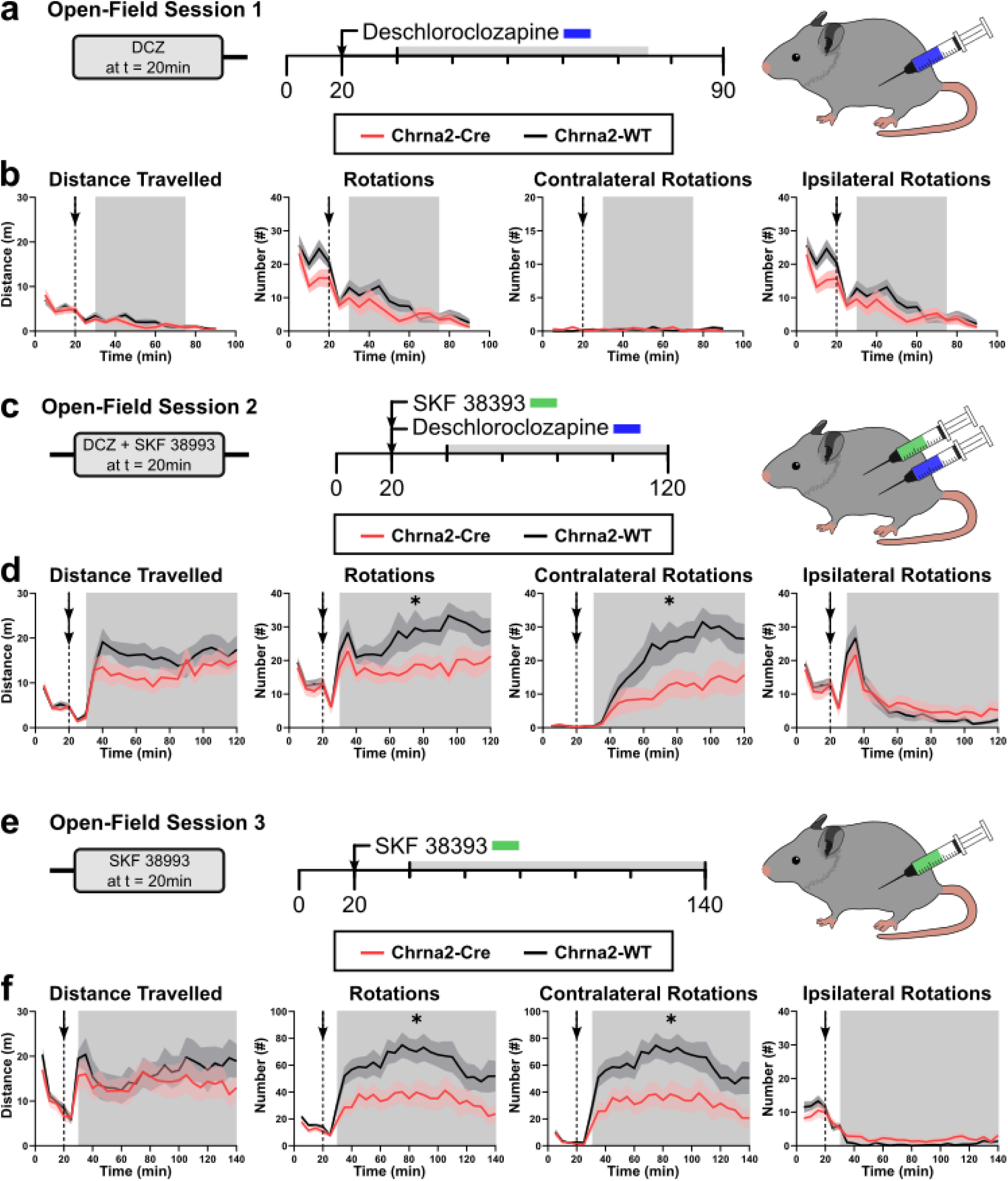
Concurrent Chrna2^+^ and D1-receptor stimulations decrease rotations in a 6-OHDA model. **a** Timeline of the first open-field session and schematic illustrating the experimental design. DCZ (100 µg/kg) was intraperitoneally administered at t = 20 min. **b** Distance travelled: genotype, F_(1,19)_ = 0.9890, p = 0.3325; time F_(8, 152)_ = 4.676, p < 0.0001; interaction, F_(8,152)_ = 1.343, p = 0.2265. Rotations: genotype, F_(1,19)_ = 0.1270, p = 0.2738; time F_(8, 152)_ = 6.655, p < 0.0001; interaction, F_(8,152)_ = 1.619, p = 0.1236. Contralateral rotations: genotype, F_(1,19)_ = 0.1456, p = 0.7070; time F_(8, 152)_ = 0.9272, p = 0.4959; interaction, F_(8,152)_ = 1.263, p = 0.2666. Ipsilateral rotations: genotype, F_(1,19)_ = 0.1290, p = 0.2701; time F_(8, 152)_ = 6.529, p < 0.0001; interaction, F_(8,152)_ = 1.263, p = 0.2666. **c** Timeline of the second open-field session and schematic illustrating the experimental design. DCZ (100 µg/kg) and SKF38393 (3 mg/kg) were intraperitoneally administered at t = 20 min. **d** Distance travelled: genotype, F_(1,19)_ = 1.997, p = 0.1738; time F_(17, 323)_ = 1.003, p = 0.4541; interaction, F_(17,323)_ = 0.5463, p = 0.9280. Rotations: genotype, F_(1,19)_ = 5.697, p = 0.0.276; time F_(17, 323)_ = 2.016, p = 0.0103; interaction, F_(17,323)_ = 0.8931, p = 0.5826. Contralateral rotations: genotype, F_(1,19)_ = 5.603, p = 0.0287; time F_(17, 323)_ = 11.00, p < 0.0001; interaction, F_(17,323)_ = 1.933, p = 0.0150. Ipsilateral rotations: genotype, F_(1,19)_ = 0.8059, p = 0.3806; time F_(17, 323)_ = 26.41, p < 0.0001; interaction, F_(17,323)_ = 1.400, p = 0.1340. **e** Timeline of the third open-field session and schematic illustrating the experimental design. SKF alone (3 mg/kg) was intraperitoneally administered at t = 20 min. **f** Distance travelled: genotype, F_(1,18)_ = 0.3713, p = 0.5499; time F_(21, 378)_ = 1.528, p = 0.0647; interaction, F_(21,378)_ = 0.8059, p = 0.7131. Rotations: genotype, F_(1,18)_ = 6.371, p = 0.0212; time F_(21, 378)_ = 4.139, p < 0.0001; interaction, F_(21,378)_ = 0.7960, p = 0.7253. Contralateral rotations: genotype, F_(1,18)_ = 6.586, p = 0.0194; time F_(21, 378)_ = 4.454, p < 0.0001; interaction, F_(21,378)_ = 0.7633, p = 0.7646. Ipsilateral rotations: genotype, F_(1,18)_ = 3.607, p = 0.0737; time F_(21, 378)_ = 2.037, p = 0.0048; interaction, F_(21,378)_ = 0.8786, p = 0.6196. The dashed line and/or arrow depict the moment the i.p. injections occurred. Statistical analysis: RM two-way ANOVA *p < 0.05, Chrna2-Cre vs. Chrna2-WT. All results are shown as mean ± SEM. n_Chrna2-Cre_ = 10, n_Chrna2-WT_ = 11. For the third open-field test session one Chrna2-WT mouse had to be removed due to non-responsiveness to the pharmacological intervention.

In conclusion, although PT activation was insufficient to counteract the motor deficits induced by unilateral dopamine loss, simultaneous Chrna2^+^ neuronal activation and SKF38393 administration resulted in hindered contralateral and overall turns of Chrna2-Cre mice (Fig. 5). This finding suggests that PT neuronal activation can selectively repress D1-receptor-driven motor output in dopamine-depleted mice.

### 3.6 Chrna2^+^ activation selectively reduced axial LID

We next examined whether repeated activation of Chrna2^+^ neurons influenced levodopa-induced dyskinesia. Chronic L-DOPA treatment in dopamine-depleted animals induces progressive sensitization of striatal circuits, leading to altered responses to subsequent interventions^38,39^. Also known as priming, it is connected with changes in neurotransmitters and receptors for dopamine and is involved in maladaptive plasticity and abnormal motor responses^39^. Following two consecutive weeks of L-DOPA priming, the animals were examined using AIM/DT ratings over the third week (Fig. 6a). No significant differences between groups were detected during the first two assessment sessions (Fig. 6b,c,e,f,h,i,k,l). However, during the final assessment session, Chrna2-Cre mice displayed a significant reduction in axial DTs (p = 0.0239), with the strongest effect observed during the fourth examination period (t = 80 min) (p = 0.0082) (Fig. 6g). No effects were observed for limb or orofacial DTs (Fig. 6j,m).

**Fig. 6:**
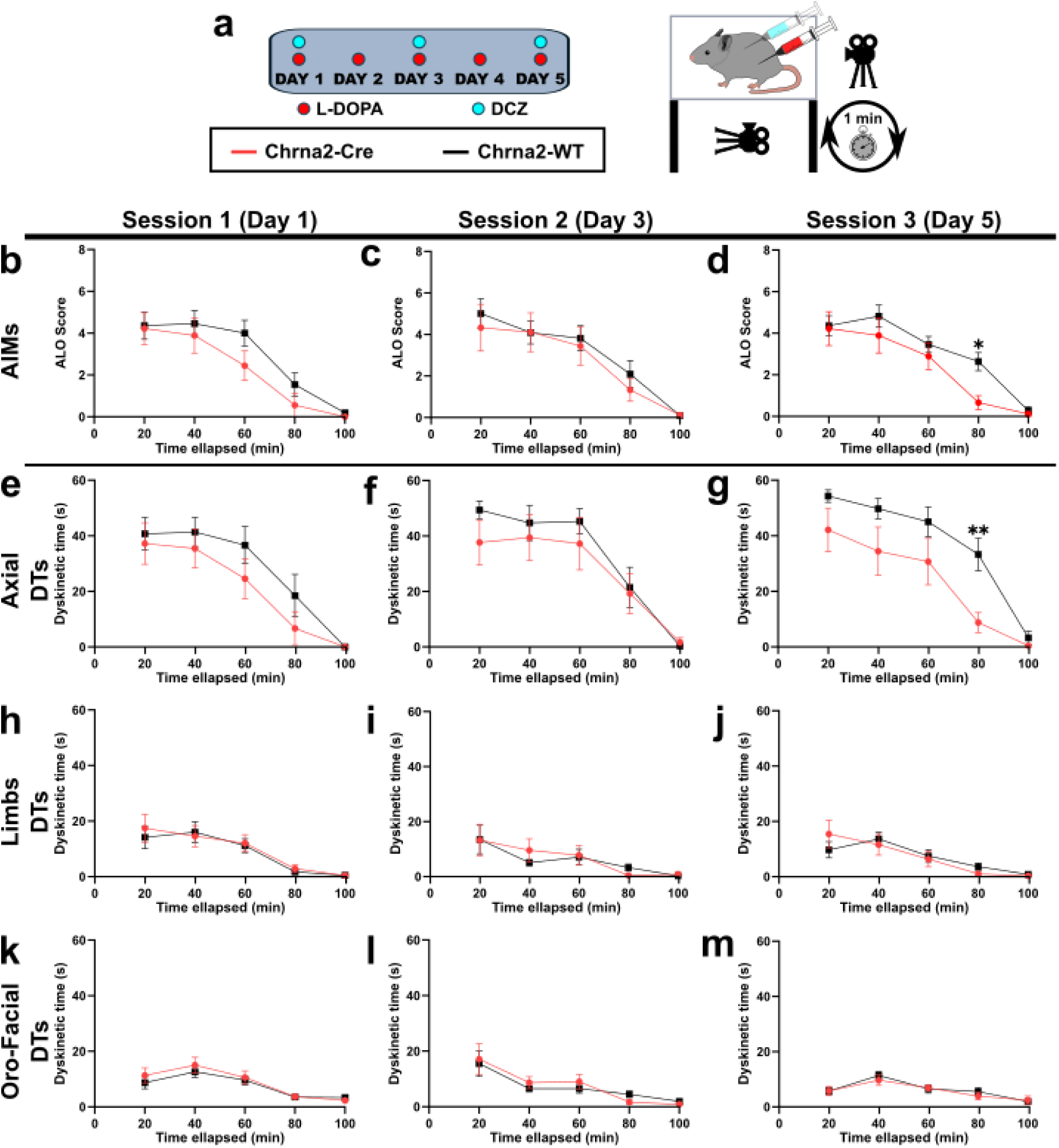
Chrna2+ activation in LID selectively reduces axial dyskinesia. **a** Timeline and schematic of dyskinesia assessment and AIMs/DTs recordings during the third week. Each animal received DCZ (100 µg/kg) and L-DOPA (6 mg/kg) on day 1, day 3, and day 5, respectively. On day 2 and day 4 (non-assessed days), the mice only received L-DOPA (6 mg/kg). **b** Session 1 AIMs: genotype, F_(1,18)_ = 1.176, p = 0.2925; time F_(4, 72)_ = 36.75, p < 0.0001; interaction, F_(4,72)_ = 0.9010, p = 0.4681. **c** Session 2 AIMs: genotype, F_(1,18)_ = 0.2429, p = 0.6280; time F_(4, 72)_ = 26.91, p < 0.0001; interaction, F_(4,72)_ = 0.2540, p = 0.9063. **d** Session 3 AIMs: genotype, F_(1,18)_ = 2.257, p = 0.1503; time F_(4, 72)_ = 37.68, p < 0.0001; interaction, F_(4,72)_ = 1.661, p = 0.1685. Šídák’s multiple comparisons test found significance at t = 80 min, *p = 0.0387. **e** Session 1 axial DTs: genotype, F_(1,18)_ = 1.016, p = 0.3268; time F_(4, 72)_ = 33.07, p < 0.0001; interaction, F_(4,72)_ = 0.7732, p = 0.5462. **f** Session 2 axial DTs: genotype, F_(1,18)_ = 0.5551, p = 0.4659; time F_(4, 72)_ = 40.19, p < 0.0001; interaction, F_(4,72)_ = 0.7414, p = 0.5669. **g** Session 3 axial DTs: genotype, F_(1,18)_ = 6.081, p = 0.0239; time F_(4, 72)_ = 44.16, p < 0.0001; interaction, F_(4,72)_ = 1.878, p = 0.1236. Šídák’s multiple comparisons test found significance at t = 80 min, **p = 0.0082. **h** Session 1 limbs DTs: genotype, F_(1,18)_ = 0.06545, p = 0.8010; time F_(4, 72)_ = 18.88, p < 0.0001; interaction, F_(4,72)_ = 0.2666, p = 0.8985. **i** Session 2 limbs DTs: genotype, F_(1,18)_ = 0.03944, p = 0.8448; time F_(4, 72)_ = 6.695, p = 0.0001; interaction, F_(4,72)_ = 0.4400, p = 0.7793. **j** Session 3 limbs DTs: genotype, F_(1,18)_ = 0.003171, p = 0.9557; time F_(4, 72)_ = 12.17, p < 0.0001; interaction, F_(4,72)_ = 1.098, p = 0.3640. **k** Session 1 oro-facial DTs: genotype, F_(1,18)_ = 0.4785, p = 0.4979; time F_(4, 72)_ = 15.67, p < 0.0001; interaction, F_(4,72)_ = 0.4150, p = 0.7973. **l** Session 2 oro-facial DTs: genotype, F_(1,18)_ = 0.04920, p = 0.8270; time F_(4, 72)_ = 11.88, p < 0.0001; interaction, F_(4,72)_ = 0.4568, p = 0.7671. **m** Session 3 oro-facial DTs: genotype, F_(1,18)_ = 0.1496, p = 0.7035; time F_(4, 72)_ = 13.26, p < 0.0001; interaction, F_(4,72)_ = 0.4324, p = 0.7847. Statistical analysis: RM two-way ANOVA *p < 0.05, **p < 0.01, Chrna2-Cre vs. Chrna2-WT. All results are shown as mean ± SEM. n_Chrna2-Cre_ = 9, n_Chrna2-WT_ = 11.

Consequently, the reduction in overall AIM scores was not statistically significant (p = 0.1503), although a significant effect was detected during the fourth observation period (t = 80 min) (p = 0.0387) (Fig. 6d). Thus, activation of Chrna2^+^ neurons selectively attenuated the axial component of LID.

### 3.7 Chrna2^+^ excitation repressed acute striatal hyperactivity in a 6-OHDA model

To investigate if our behavioural findings can be traced to changes in underlying circuit mechanisms, we quantified both chronic (ΔFosB) and acute (c-Fos) striatal neuronal activation upon dopaminergic agonism. Opposingly to c-Fos, whose half-life is of approximately one hour^40^, ΔFosB retains its levels for weeks after the last L-DOPA administration, allowing us to monitor how LID behaviour correlates with fluctuations in striatal activity^13,15^. As expected, both genotypes displayed robust increases in ΔFosB^+^ exclusively in the lesioned striatum (Chrna2-Cre – p < 0.0001, Chrna2-WT – p < 0.0001). However, no significant effect was detected when comparing Chrna2-Cre and Chrna2-WT corresponding striata (p = 0.4029), indicating that repeated Chrna2^+^ activation did not alter chronic LID-associated ΔFosB accumulation (Fig. 7a-c).

**Fig. 7:**
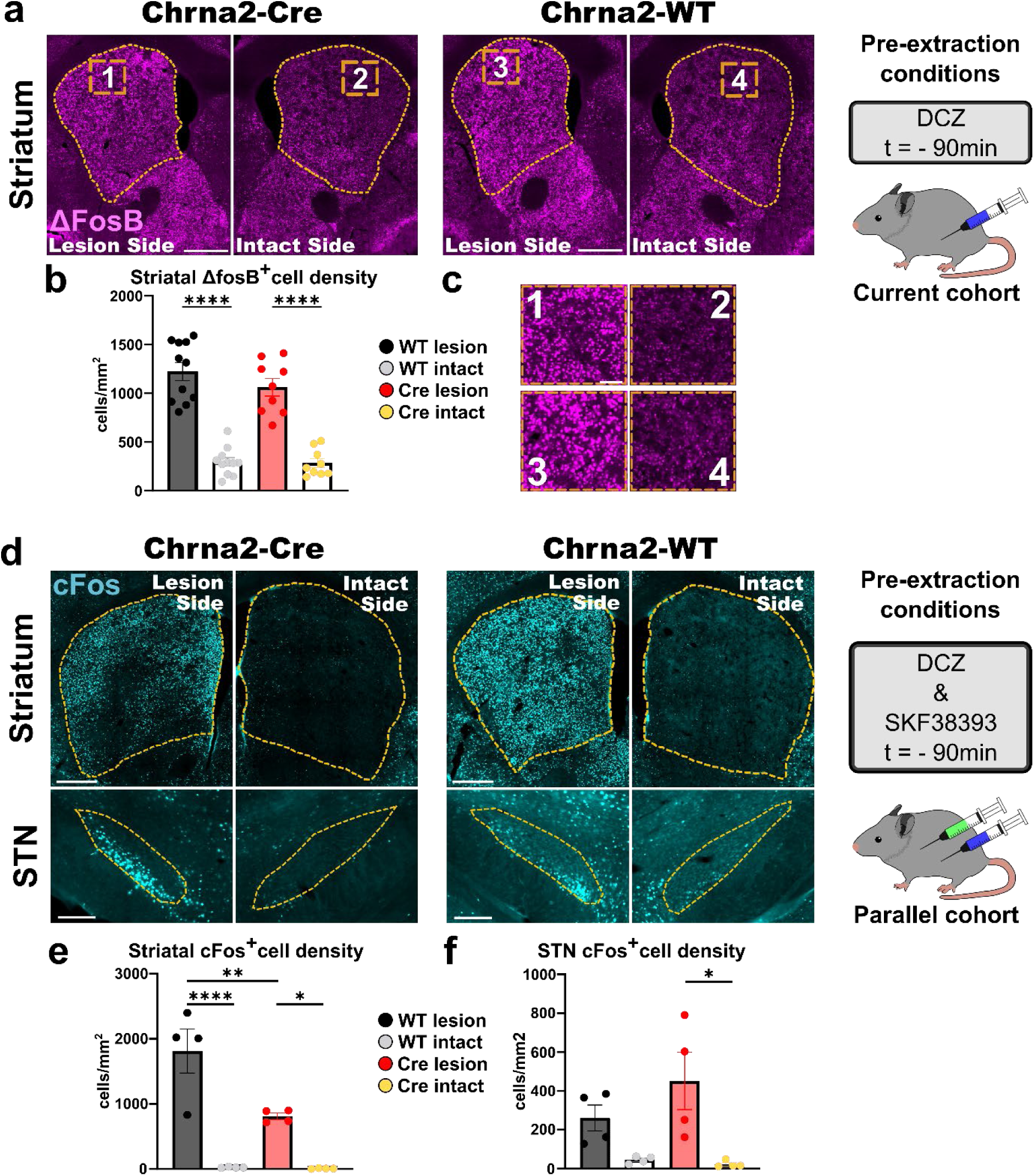
Chrna2^+^ activation repressed acute, but not chronic striatal neuronal excitation in a 6-OHDA model. **a** Representative fluorescent images depicting intact and striatal chronic neuronal activation (ΔFosB, violet) in Chrna2-Cre and Chrna2-WT animals in 6-OHDA-lesioned mice. Scale bar: 500 µm. **b** Striatal ΔFosB^+^ cell density analysis: F_(3, 36)_ = 5.127, p < 0.0001. Chrna2-Cre lesion vs. Chrna2-Cre intact: p < 0.0001, Chrna2-WT lesion vs. Chrna2-WT intact: p < 0.0001, Chrna2-Cre lesion vs. Chrna2-WT lesion: p = 0.4029, Chrna2-Cre intact vs. Chrna2-WT intact: p = 0.9994. **c** Zoomed-in images highlighting cellular mapping of ΔFosB expression. Scale bar: 50 µm. **d** Representative fluorescent images depicting intact and ipsilateral acute neuronal activation (c-Fos, turquoise) in the dorsal striatum (upper row) and STN (bottom row) in Chrna2-Cre and Chrna2-WT in 6-OHDA-lesioned mice. Dorsal striatum scale bar: 500 µm, STN scale bar: 100 µm. **e** Striatal c-Fos^+^ cell density analysis: F_(3, 12)_ = 1.810, p < 0.0001. Chrna2-Cre lesion vs. Chrna2-Cre intact: p = 0.0278, Chrna2-WT lesion vs. Chrna2-WT intact: p < 0.0001, Chrna2-Cre lesion vs. Chrna2-WT lesion p = 0.0065, Chrna2-Cre intact vs. Chrna2-WT intact: p = 0.9999. **f** STN c-Fos^+^ cell density analysis: F_(3,12)_ = 22.67, p = 0.0088. Chrna2-Cre lesion vs. Chrna2-Cre intact: p = 0.0131, Chrna2-WT lesion vs. Chrna2-WT intact: p = 0.2891, Chrna2-Cre lesion vs. Chrna2-WT lesion p = 0.3811, Chrna2-Cre intact vs. Chrna2-WT intact: p = 0.9973. Statistical analysis: one-way ANOVA *p < 0.05, **p < 0.01, ***p < 0.001, ****p < 0.0001, Chrna2-WT lesion vs. Chrna2-WT intact vs. Chrna2-Cre lesion vs. Chrna2-Cre intact for c-Fos and ΔFosB, followed by Tukey’s multiple-comparisons test. All results are shown as mean ± SEM. ΔFosB analysis: n_Chrna2-Cre_ = 9, n_Chrna2-WT_ = 11, c-Fos and enkephalin analyses: n_Chrna2-Cre_ = 4, n_Chrna2-WT_ = 4. Any cell unable to qualify as activated was excluded from the analysis.

We then investigated acute neuronal activity patterns (c-Fos)^27^ of the 6-OHDA-lesioned histology cohort (Methods 2.2.3) upon acute stimulation with DCZ (100 µg/kg) and SKF38393 (3 mg/kg) in the striatum and STN. Although c-Fos levels remained elevated in the 6-OHDA-lesioned striata of both genotypes (both p < 0.0001), Chrna2-Cre mice displayed significantly lower striatal c-Fos expression than -WT controls (p = 0.0065), indicating suppression of acute striatal overactivity (Fig. 7d,e). Regarding the STN, Chrna2^+^ activation did not have any significant effect upon c-Fos levels when comparing complementary regions of 6-OHDA-lesioned Chrna2-WT and -Cre (lesioned-lesioned: p = 0.3811, intact-intact: p = 0.9973) (Fig. 7d,f), albeit an interhemispherical significance in Chrna2-Cre mice (p = 0.0131) (Fig. 7d,f).

Lastly, c-Fos/enkephalin co-labelling demonstrated that a marginal number of activated striatal neurons expressed the D2-SPN-selective marker enkephalin, indicating that most striatal activity induced by SKF38393 originated from D1-SPNs rather than D2-SPNs (Fig. 8). Collectively, these findings suggest that PT activation selectively dampens acute striatal hyperactivity associated with D1-SPN stimulation, without affecting ΔFosB-associated maladaptive plasticity from previous chronic L-DOPA treatments.

**Fig. 8:**
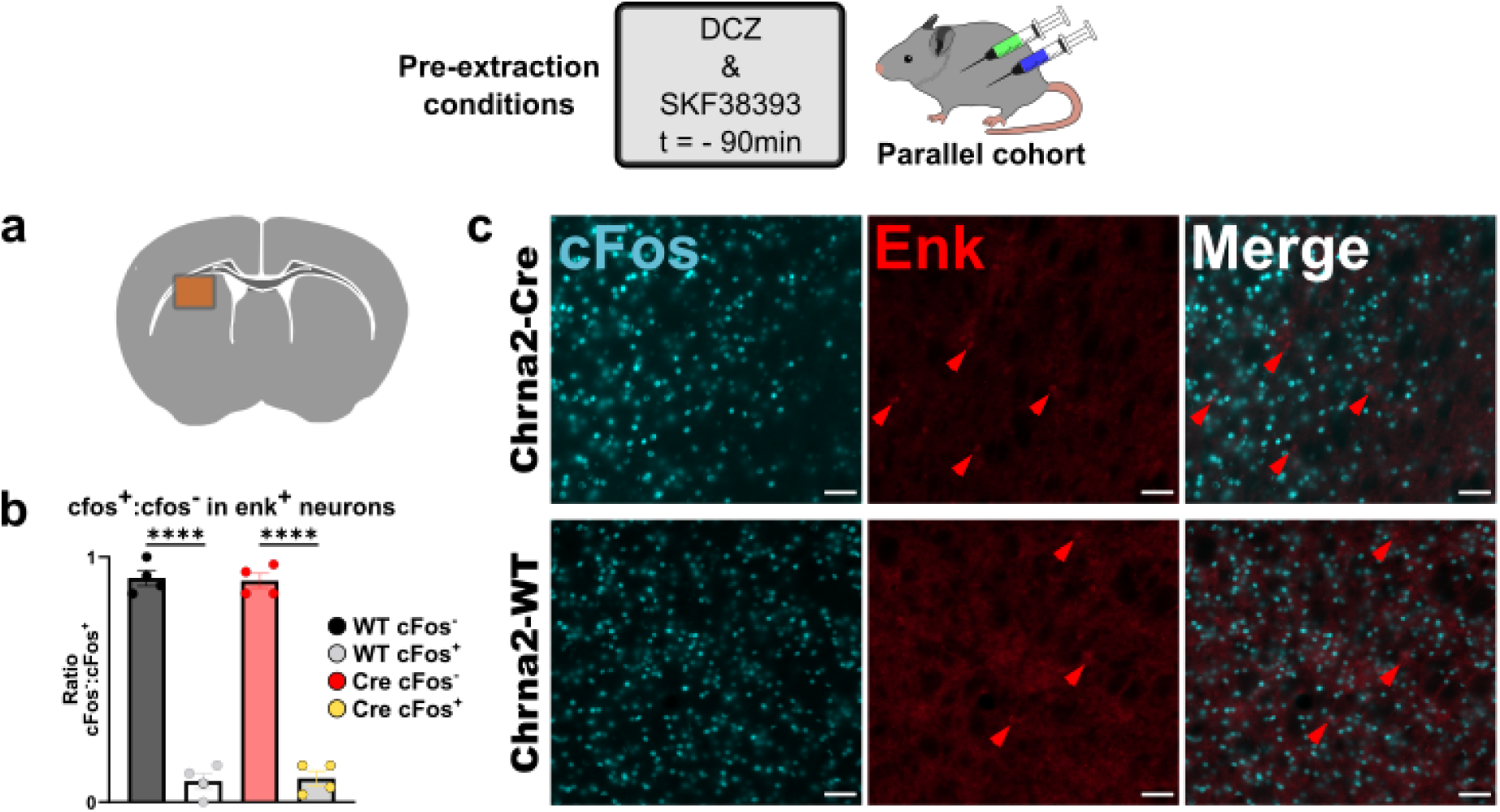
Acute striatal activation primarily stems from non-D2-SPNs. **a** Representative brain slice and ROI (rectangle) of a quantification area. **b** Zoomed-in pictures from the coronal brain slice showing enkephalin (enk, red), c-Fos (blue), and merged channels for Chrna2-Cre and Chrna2-WT lesioned dorso-lateral striatum. Scale bar 50µm. **c** Enkephalin positive (enk^+^) striatal cells with and without c-Fos signal (c-Fos^+^/c-Fos^-^) in Chrna2-Cre and Chrna2-WT lesioned striatum. F_(3, 12)_ = 0.02911, p < 0.0001. c-Fos^-^Chrna2-WT vs. c-Fos^+^ Chrna2-WT p < 0.0001, c-Fos^-^Chrna2-Cre vs. c-Fos^+^ Chrna2-Cre p < 0.0001, c-Fos^-^Chrna2-WT vs. c-Fos^-^Chrna2-Cre: p > 0.9999, c-Fos^+^ Chrna2-WT vs. c-Fos^+^ Chrna2-Cre p > 0.9999. Statistical analysis: one-way ANOVA *p < 0.05, **p < 0.01, ***p < 0.001, ****p < 0.0001, n_Chrna2-Cre_ = 4, n_Chrna2-WT_ = 4. c-Fos^-^Chrna2-WT vs. c-Fos^+^ Chrna2-Cre vs. c-Fos^+^ Chrna2-WT vs. c-Fos^-^ Chrna2-Cre for enkephalin^+^ graph, followed by Šídák’s multiple comparisons test. All results are shown as mean ± SEM. Any cell unable to qualify as activated was excluded from the analysis.

## 4. Discussion

PT neurons represent a major cortical output pathway to subcortical motor circuits, yet their contribution to pathological motor states remains poorly addressed. In the present study, we examined the behavioural and neurobiological consequences of chemogenetic activation of motor-cortical PT neurons in both intact and dopamine-denervated mice. In healthy animals, unilateral PT activation suppressed spontaneous locomotor activity in the open field, indicating a net motor-inhibitory effect of enhanced PT output. In dopamine-depleted mice, this activation attenuated D1-receptor-mediated rotational behaviours, reduced axial levodopa-induced dyskinesia, and diminished acute D1-dependent striatal activation. To the best of our knowledge, our investigation is the first to reveal functional effects of a motor-cortical PT stimulation in a rodent model of Parkinson’s disease. The findings identify PT neurons as modulators of pathological dopaminergic motor states and suggest a previously underappreciated interaction between cortical PT output and striatal dopamine signalling. Collectively, these findings reveal a functional cortical-striatal interaction, whereby increased PT output can mitigate specific manifestations of dopamine-dependent motor dysfunction, providing new insight into the circuit mechanisms that shape parkinsonian motor behaviour and dyskinesia.

### 4.1 PT activation reduces locomotion in healthy mice and contraversive turns in a 6-OHDA model

It has been shown that PT neurons furnish essential inputs to the striatum and other deeper brain regions and are therefore critically positioned to influence movement execution and control^1,41,42^. PT chemogenetic activation encumbered locomotion; the net physiological summation of their descending, innervated brain areas culminated in a hypokinetic phenotype. It is important to reiterate that our study encompasses an *in toto* activation of PT neurons, rather than employing pathway-selective approaches^43,44^. The absence of a prokinetic effect contrasts with previous studies reporting improved locomotion following optogenetic stimulation of cortico-fugal neurons^44,45^. In our study, we find no general motor improvements upon PT chemogenetic stimulation alone. Differences in cortical area, neuronal population, projection target, and stimulation paradigm may critically shape the behavioural consequences of cortical motor circuit activation. Together, these findings propose that the effects of cortico-fugal stimulation are highly circuit-specific and cannot be generalized across cortical output pathways.

Because PT activation alone did not improve locomotion in 6-OHDA-lesioned mice, we reasoned that its functional impact might become more legible under an enhanced dopamine D1-receptor drive. To test this hypothesis, we administered the D1-receptor agonist SKF38393, known for inducing contralateral rotations and locomotor hyperactivity in unilateral 6-OHDA-lesioned mice^34,37^. Notably, dopamine depletion in mice provokes an ipsiversive rotational bias towards the striatum with less signalling outputs^34,36^. Accordingly, previous literature^15,18,46–48^ has illustrated that discrete and unilateral modulations of striatal neurons lead to either ipsi-or contraversive rotational preferences. Furthermore, dopamine D1-receptors expressed on striatonigral neurons become hypersensitised to dopamine agonism^34,36,49^ (i.e., SKF38393). Against this background, PT activation attenuated SKF38393-induced rotational behavior. Congruently, the functional impact of simultaneous dopaminergic and PT stimulations revealed a state-dependent interaction between cortical output and dopamine signalling. The attenuation of turning behaviour therefore suggests that an increased PT output counteracts excessive D1-mediated motor drive in the dopamine-depleted striatum, though this interaction remains to be established.

Notably, the attenuating effect on D1-receptor simulation induced rotational behaviour persisted during subsequent testing sessions. This sustained effect raises the possibility that transient DCZ-mediated PT activation induced longer-lasting changes within cortico-striatal circuits. Although the underlying mechanisms remain unclear, DCZ exhibits high DREADD affinity, rapid brain penetration, and minimal off-target effects compared with earlier chemogenetic ligands^30,50,51^. Whether repeated PT activation can engage adaptive or maladaptive plasticity within motor circuits remains unknown and warrants further investigation.

Our behavioural data strongly suggest that a dopaminergic stimulation is necessary to mediate motor restoration or improve motility at a minimum, conversely to the findings of a previous investigation^52^ arguing that dopamine is expandable for movement initiation. The rotational outputs of ensuing PT chemogenetic activation indicate a strong striatal implication^53^. Future studies that combine cell-type-specific manipulations with longitudinal circuit recordings could determine whether PT neurons exert lasting effects on the functionality of their cortico-striatal network.

### 4.2 PT neurons selectively dampen axial dyskinesia

A body of experimental literature has established a link between LID and hypersensitization of striatal dopamine D1-receptors^38,39^. Although a faulty activity balance between D1- and D2-SPNs forms its fundamental mechanism, the pathophysiology of LID is complex and involves several neural pathways^54^. In our AIM/DT scoring design, PT activation selectively reduced axial LID. While PT activation did not affect limb or oro-facial dyskinesia, its impact on axial domains indicates that cortical output pathways may regulate specific dyskinetic motor domains. Observations in recent studies emphasise that to alleviate LID effects, a co-activation of D1- and D2-SPNs is needed, and that activating D2-SPNs would alleviate LID^53,54^. Deconstructing the sophisticated D1- and D2-SPNs systems, a synchronous coexistence is required between the action-facilitating former and competing action-suppressing latter^12^ to ensure precise action selection and movement generation. Our hypothesis that PT neurons have a therapeutic effect on axial dyskinesia resonates with the theory stating that as dyskinesia materialises upon L-DOPA administration, D1-SPNs heightened activity persists, while D2-SPNs’ is noticeably repressed^54^. Recent studies in LID models^18^ have demonstrated a strong dependence of axial dyskinesia from D2-SPNs^55^. In our case, a tonic PT stimulation counteracts L-DOPA-induced abnormal axial kinetics in experimental parkinsonism through a possible restoration of the D2:D1 SPNs balance. Therefore, PT neurons may qualify as candidates of a previously unrecognised cortical node, capable of modulating clinically relevant aspects of dyskinetic motor behaviour. Lastly, the DTs method developed by our group in this investigation offers a more precise and sensitive real-time add-on to the widely validated AIM rating methods used in dyskinetic research^15,18,19,53,56^.

### 4.3 PT activation dampens striatal hyperactivity in a 6-OHDA model

Based on previous behavioural results on acute and chronic dopaminergic treatments, we assessed whether these could be correlated to changes in striatal activity. Acute D1-receptor stimulation with SKF38393 induced robust c-Fos expression in the dopamine-depleted, hypersensitised striatum. Notably, chemogenetic PT activation significantly reduced c-Fos levels, echoing the attenuation of D1-receptor-mediated rotational behaviour. Contrastingly, this chemogenetic intervention did not impact accrued striatal ΔFosB levels, induced by a chronic levodopa regimen.

Both c-Fos and ΔFosB are established markers of dopamine-dependent striatal activation in 6-OHDA models^40,57^. Whereas c-Fos reflects acute neuronal activation, ΔFosB accumulates following repeated dopaminergic stimulation and has been closely linked to the development and severity of levodopa-induced dyskinesia^57,58^. Following three weeks of L-DOPA administration, all mice demonstrated elevated ΔFosB levels in the lesioned striatum. The aggregation of this transcription factor could have potentially overridden the less frequent PT activations that led to an alleviation of LID^57,59^. In 6-OHDA neurotoxin models, L-DOPA administration shifts the D1-vs. D2-SPNs activity balance, exacerbating the former while dampening the latter^60^. Previously, L-DOPA-driven ΔFosB levels have been primarily observed in D1-SPNs^61^. Importantly, reduced c-Fos expression closely mirrored the attenuation of contraversive turning behaviours, supporting a functional liaison between PT activation and suppression of excessive striatal output. The observed shift in rotations and striatal activity follows the link between hemispheric differences of striatal output, directly correlating with rotational bias^33,56^. In contrast, we detected no significant changes in subthalamic nucleus activation, suggesting that the behavioral effects were more closely associated with altered striatal processing than with gross changes in STN recruitment.

The cellular mechanisms underlying these effects remain uncertain. Our histological analyses revealed only limited c-Fos expression in enkephalin^+^, dopamine D2-receptor-expressing neurons, suggesting that the observed changes may predominantly involve D1-SPNs. The c-Fos levels in enkephalin^-^cell populations could be linked to inhibitory influences via cholinergic interneurons from other striatal subdomains^8^. Echoing with previous hypotheses arguing for preferential striatal D1-SPN connection^62^ and consequent to our behavioural readouts, we suggest that a PT chemogenetic stimulation may indirectly recruit striatal D1-SPNs in a 6-OHDA lesion model of PD, thus stabilising turning behaviours and striatal activity balance. This conjecture is supported by previous investigations that dissect the cortico-striatal system, illustrating PT affinity towards D1-SPNs^62^. Although other studies support a D2-SPNs-favored PT innervation in naïve systems^63,64^, our analysis yielded minimal D2-SPNs activation.

However, the present data do not attribute striatal excitation to a discrete neuronal subset. PT neurons directly innervate SPNs and can regulate striatal processing through local interneuronal networks^8,63–68^. Consequently, the observed c-Fos reduction may have resulted from multiple circuit mechanisms, rather than being caused by an individual neuronal population. Converging behavioural and molecular findings suggest that PT activation influences D1-receptor signalling in a dopamine-denervated state.

## Conclusion

Current models of Parkinson’s disease largely focus on dopamine depletion and downstream basal ganglia dysfunction. Our findings extend this framework by identifying cortical PT neurons as active modulators of pathological dopaminergic motor responses. Rather than directly restoring movement, PT activation selectively limited maladaptive responses to dopaminergic stimulation (i.e., turning behaviours and axial dyskinesia). We speculate that PT stimulation engages striatal microcircuitry to restore the equilibrium between D1-SPNs and D2-SPNs activity^8,67,68^. Future studies combining cell-type-specific manipulations with electrophysiological and calcium-imaging approaches would be needed to define the striatal subpopulations and circuits PT activation engage. In sum, our results reveal a previously unapprised role for cortical output pathways in regulating pathological motor states and suggest that PT neurons represent a promising substrate for future circuit-based therapies. More broadly, the study highlights the importance of considering cortico-striatal interactions when developing or improving neuromodulation strategies for Parkinson’s disease^10,69,70^.

## Acknowledgments

We thank Sascha Gude for animal handling at the Hvidovre Copenhagen University Hospital and the Core Facility for Integrated BioImaging at the University of Copenhagen for their support. We thank Osama Elabi, Markus Aldén, and Teodor Nyman for their valuable help and advice in setting up the mouse LID model.

## Data availability

A source data file supporting the findings of this study will be provided for this paper. Further data will be made available upon request.

## Author Contributions

**Mihai Atudorei** (MA):

investigation, formal analysis, data curation, visualization, writing – original draft, editing

**Gustaf Olsson** (GO):

investigation, formal analysis, data curation, visualization, writing – original draft, editing

**Christian del Agua Villa** (CAV):

investigation, formal analysis, data curation

**Claudiu Corcea** (CC):

investigation

**Ulrik Gether** (UG):

resources, editing

**Maria Angela Cenci** (MAC):

conceptualization, supervision, resources, funding acquisition, editing

**Hartwig Roman Siebner** (HS):

conceptualization, supervision, funding acquisition, writing – original draft, editing, project administration

**Mattias Rickhag** (MR):

conceptualization, supervision, funding acquisition, writing – original draft, editing, data curation, formal analysis, project administration

